# A combinatorial approach for the synthesis of multi-phosphorylated peptides: new tool for studying phosphorylation patterns in proteins

**DOI:** 10.1101/196444

**Authors:** Mamidi Samarasimhareddy, Daniel Mayer, Norman Metanis, Dmitry Veprintsev, Mattan Hurevich, Assaf Friedler

## Abstract

Phosphorylation of proteins at multiple sites creates different phosphorylation patterns that are essential for their biological activity. For example, such patterns contribute to the redirection of signalling to alternative pathways. Multi-phosphorylated peptides are excellent tools to systematically study the impact of unique phosphorylation patterns on signalling, but their synthesis is extremely difficult. Here we present an efficient and general method for the synthesis of multi-phosphorylated peptides, using a combination of different tailor-made coupling conditions. The method was demonstrated for the synthesis of a library of Rhodopsin C terminal peptides with distinct phosphorylation patterns containing up to seven phosphorylated Ser (pSer) and Thr (pThr) residues in close proximity to one another. Our method can be used to synthesize peptides incorporating multiple phosphorylated amino acids at high efficiency. It does not require any special expertise and can be performed in any standard peptide laboratory. This approach opens the way for quantitative mechanistic studies of phosphorylation patterns and their biological roles.

## Introduction

Phosphorylation plays a vital role in the regulation of cellular processes and is a hallmark of signal transduction.^1-3^ It can regulate the activity of proteins in many different ways, for example by inducing conformational changes or modulating interactions with other biomolecules.^4,5^ Various cell-cycle regulatory proteins and proteins involved in signalling show phosphorylations at multiple sites, which may result in a distinct signalling outputs.^6,7^ A number of studies highlighted the importance of the phosphorylation patterns for protein activities such as DNA binding, protein-protein interaction, nuclear import and export, receptor activation and many other functions.^8-10^

To elucidate the role of each individual phosphorylation pattern in signaling, multi-phosphorylated peptides with close to 100% homogeneity should be obtained. Enzymatic phosphorylation can rarely provide this due to the low specificity of kinases and extensive purification steps that are required afterwards (Fig. 1A).^11^ Chemical peptide synthesis allows site-specific phosphorylation and provides an excellent method to obtain homogeneous phosphopeptides (Fig.1B).^12,13^ However,this is a very difficult approach and only a few successful examples were reported.^14-17^ To carry out studies of the biological significance and roles of phosphorylation patterns feasible, there is a major unfulfilled need for a general synthetic method to access a library of multi-phosphorylated peptides at a low cost, high purity and sufficient yield. An efficient synthetic strategy will allow the community to address the exact biological role of each phosphorylation site/pattern of the parent protein.

**Figure 1.**
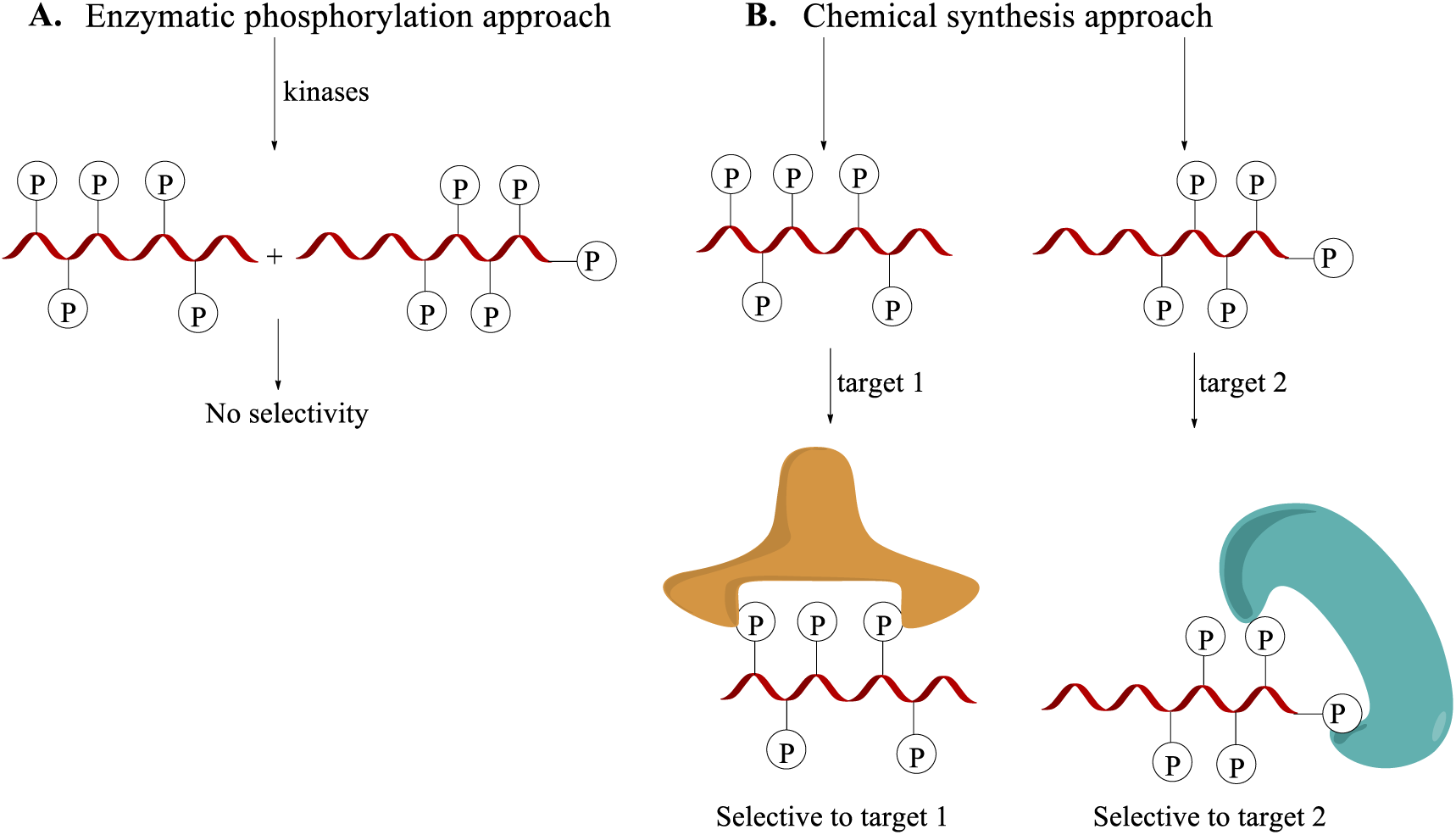
(A) Enzymatic phosphorylation is not specific or selective, leading to a mixture of peptides with different phosphorylation patterns. (B) Chemical peptide synthesis allows the preparation of multi-phosphorylated peptides with specific required patterns and can be used to elucidate the function of each modification.

While single phosphorylated peptides can be synthesized efficiently by employing standard SPPS protocols, the synthesis of multi-phosphorylated peptides using traditional methods remains extremely difficult due to steric hindrance that decreases the coupling yield, the lack of suitable protecting group strategies and incompatible cleavage conditions.^14,17-19^ The Boc-SPPS approach, which found some success, have been mostly abandoned due to the extremely hazardous HF cleavage conditions and its incompatibility with the building blocks.^20,21^

The synthesis of several multi-phosphorylated peptides that employ conventional Fmoc-SPPS strategy was achieved by following laborious coupling procedures^22^ and large excess of commercially available protected pAAs (phosphorylated amino acids) and repetitive coupling cycles.^12,14,23-26^ Microwave (MW) assisted Fmoc-SPPS of phosphorylated peptides was utilized to increase the coupling efficiency but still employs large excess pAAs and was demonstrated only for peptides in which the pAAs are not in close proximity.^27,28^

The synthesis of peptides bearing multiple and adjacent pThr residues is even more difficult than the synthesis of peptides with well separated phosphorylated amino acids or containing only pSer amino acids. Moreover, even with the use of MW assisted Fmoc-SPPS, the successful synthesis of peptides bearing multiple and adjacent pThr residues was never reported.^17,24^ Most reports describing the MW assisted SPPS of multi-phosphorylated peptides use identical coupling conditions for the entire synthesis (Fig. 2A). We found that such strategy is highly inefficient for peptides bearing multiple pThr moieties or when the pAAs are in close proximity (Fig. 2A). Extended coupling times and a large excess of highly expensive reagents and solvents may be used for the synthesis of one multi-phosphorylated peptide in a very small quantity, but this is not viable for the synthesis of a library of multi-phosphorylated peptides.

**Figure 2.**
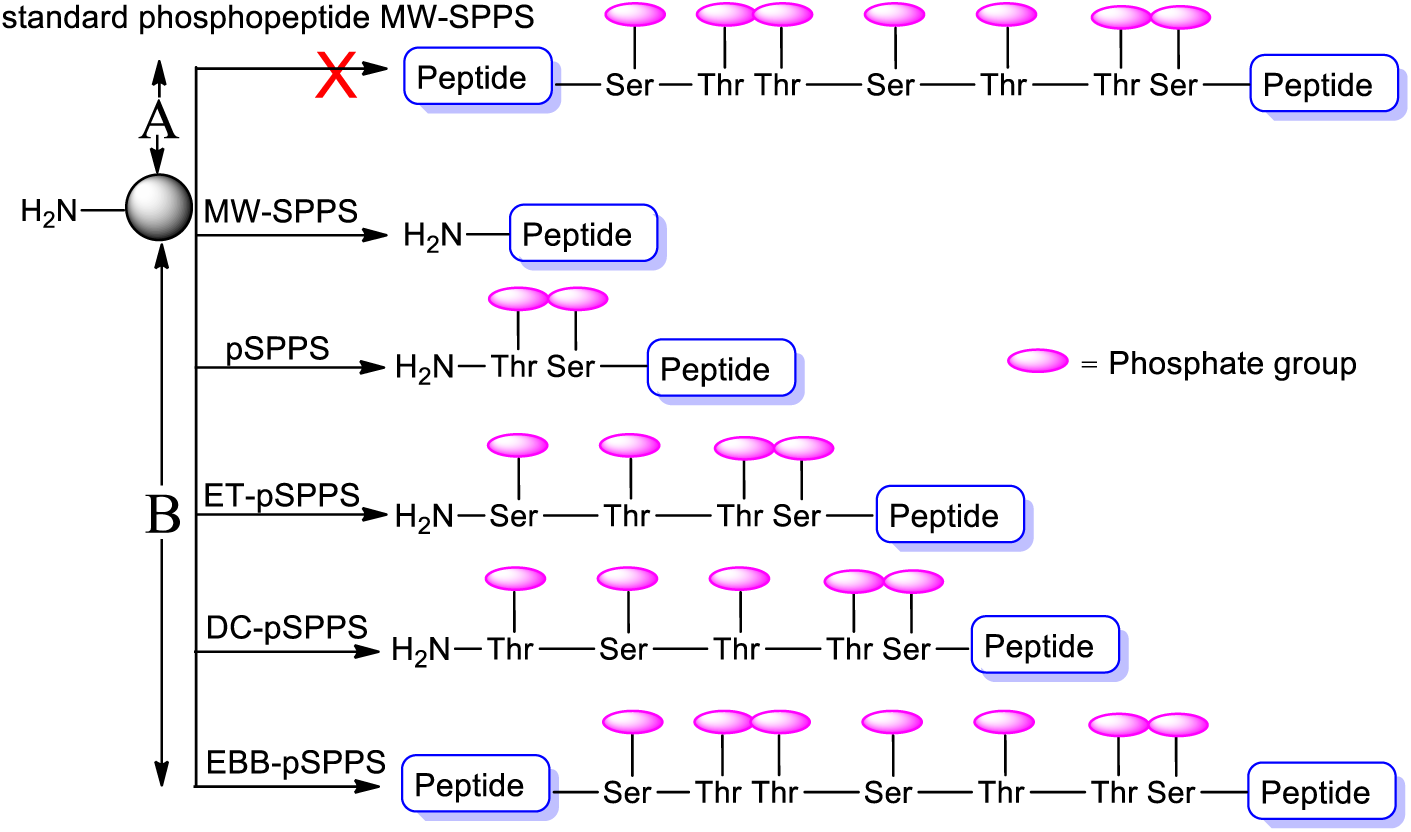
Strategies for the synthesis of multi-phosphorylated peptides. A) <u>The commonly used strategy</u>: A <u>standard phosphopeptide MW-SPPS</u> method is used throughout the entire synthesis. B) <u>Our new strategy</u>: A combination of coupling methods were used according to the specific phosphorylation pattern. ^*^coupling methods used: MW-solid phase peptide synthesis (SPPS), phospho-SPPS (pSPPS), extended coupling time pSPPS (ET-pSPPS), double coupling-pSPPS (DC-pSPPS) and excess building block pSPPS (EBB-pSPPS).

Here we present a combinatorial strategy for the MW assisted Fmoc-SPPS of highly difficult to synthesize multi-phosphorylated peptides. Our strategy employs the combination of different coupling methods by changing the coupling time, equivalents of pAAs and the repetitive coupling cycles (Fig. 2). The coupling methods are modified only when the simpler method cannot be used since it is not efficient enough for that particular coupling (Fig. 2B). The overall strategy applied here will make the approach more efficient and economical using reduced amounts of the expensive pAAs in comparison with the standard MW-phosphopeptide synthesis. We demonstrate that the synthesis of even the most hindered multi-phosphorylated peptides can be achieved in an efficient and cost-effective way by the simple modification of the coupling method.

Our model peptide for this study was derived from the C-tail of rhodopsin (Rho), a light sensitive G protein-coupled receptor (GPCR).^29-31^ The C-terminal domain of rhodopsin protein (Rho330-348, DDEASTTVSKTETSQVAPA) becomes highly phosphorylated at Ser and Thr residues (underlined) during light activation, leading to an increased affinity of arrestin towards the receptor and inhibiting further G protein signalling.^32-34^

Synthetic and homogeneous multi-phosphorylated peptides derived from this domain are essential tools to study the specific impact of each phosphorylation site or pattern on the regulatory function of rhodopsin.

Rho(330-348) contains a unique combination of four pThr and three pSer residues in very close proximities, making it very difficult to synthesize. Hence this peptide was used as a model for developing the combinatorial strategy for the synthesis of a library of multi-phosphorylated peptides in a very efficient manner. To make the strategy general and easily accessible we relied on commonly used reagents and methods with the aim to develop a method that provides efficient coupling yields without using large amounts of the very expensive pAAs. We used several coupling methods to adjust the coupling conditions to each phosphorylation pattern. The strategy developed here is very simple and can be performed in any standard peptide laboratory, enabling the synthesis of multi phosphorylated peptides with a minimal use of protected pAAs.

## Results and discussion

The coupling was performed using commercially available Fmoc-Ser(HPO_3_Bzl)-OH and Fmoc-Thr(HPO_3_Bzl)-OH and MW assisted coupling at 75°C. The coupling was performed by applying a microwave power of 25W and not allowing the temperature to reach 75°C.^35^ Fmoc deprotection was performed at room temperature to avoid β-elimination side reaction^23,28,36^. We selected the standard and widely used activating system HATU/DIEA since our aim was to make the procedure as easy and common as possible.^24^ Initially, the synthesis of peptide **1** was attempted by employing a standard MW assisted phosphopeptide coupling method. We observed a dramatic decrease in yield after the insertion of the third and the fourth pAAs (Fig. S1, S2). Peptide **1** includes seven phosphorylation sites within a ten amino acids sequence, four of which are pThr. This indicates the complexity of the sequence (Fig. 3). Since the standard MW assisted phosphopeptide coupling method did not work for this type of peptide, the coupling conditions for each amino acid were evaluated for the synthesis of peptide **1**. These included the reaction time, the amount of pAAs and the number of coupling cycles.

**Figure 3.**
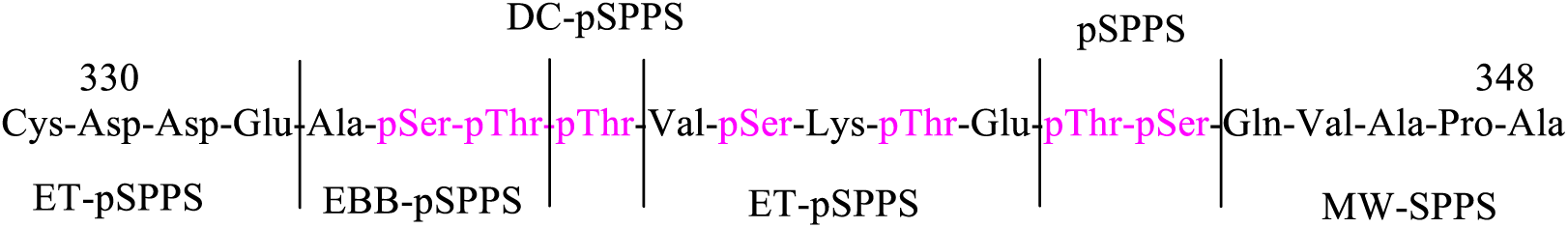
Combination of coupling methods used for the synthesis of peptide **1** (the phosphorylation sites are highlighted in pink colour).

When the existing condition was not working efficiently, we modified the coupling method by changing the time, coupling cycle or amount of pAAs (Fig. 3). Only one coupling parameter (either the number of equivalents of pAA, coupling time or number of coupling cycles) was modified each time. Using this strategy, optimized tailor made conditions were applied for each pAA in peptide **1**. The coupling methods were modified at every stage of the synthesis in accordance with the number of pAAs and the type of the pAA (See Fig. 3 and Table 1 for the details of the coupling methods). Each coupling of the of pAAs proceeded to more than 99% completion as monitored by Kaiser-ninhydrin test^37^ and by performing analytical RP-HPLC and MS analysis of the crude peptide.

**Table 1.**
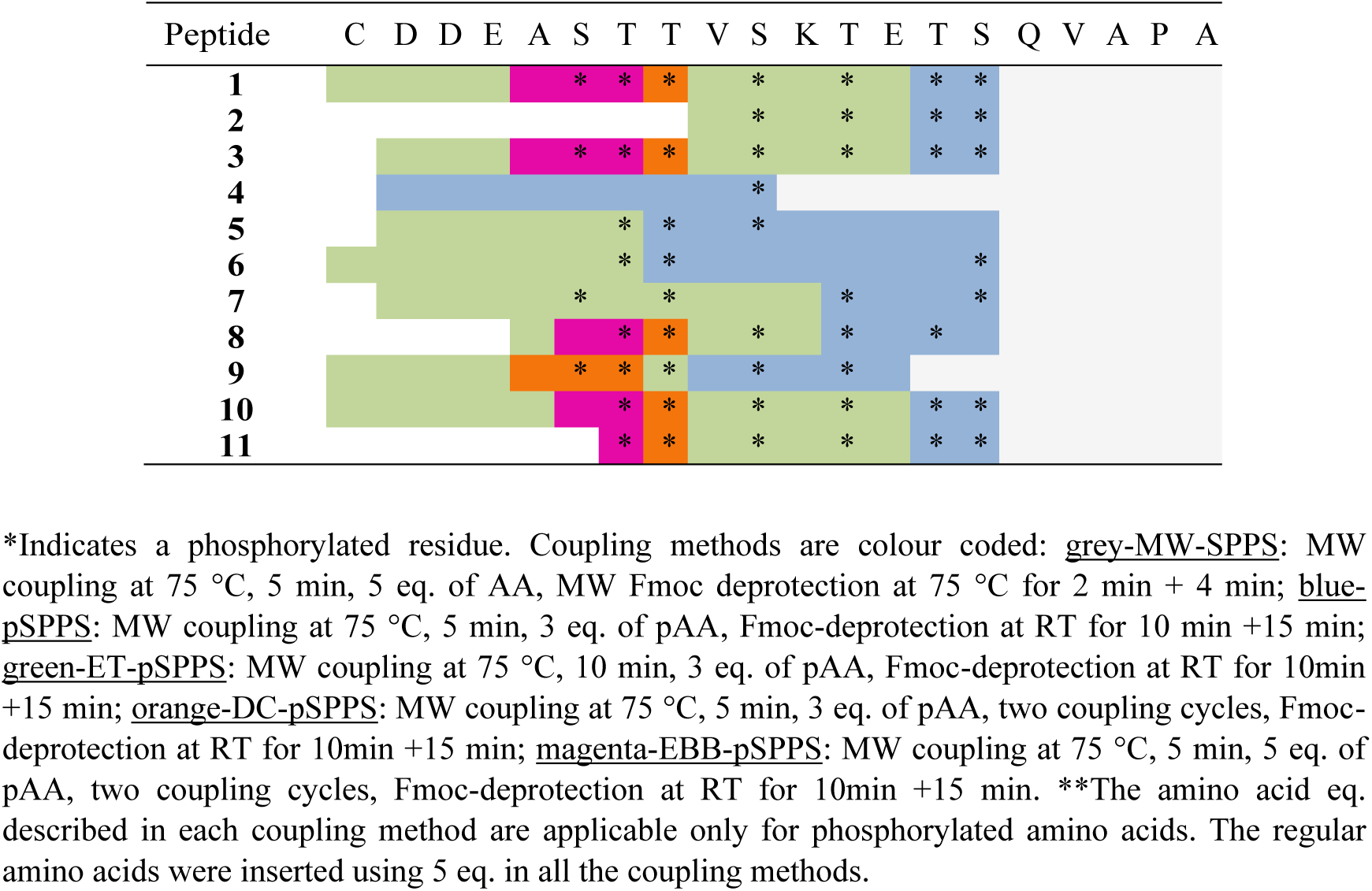
Coupling methods used for the insertion of each specific pAA in to the peptide library.

Table 1 summarizes the coupling conditions used for the insertion of each amino acid into peptide **1** (see SI Fig. S4-S6 and Table S1 for the full synthesis details of peptide **1**). Employing combinatorial approach by changing the coupling methods in correlation with the type, proximity and location of the pAA provided a highly pure multi-phosphorylated peptide **1** (Fig. 4A) and was further confirmed by analytical RP-HPLC, ESI-MS (Fig. 4B and 4C). ^31^P NMR was also recorded to determine the presence of seven phosphorylation sites in peptide **1**(Fig. 4D and Fig. S43).

**Figure 4.**
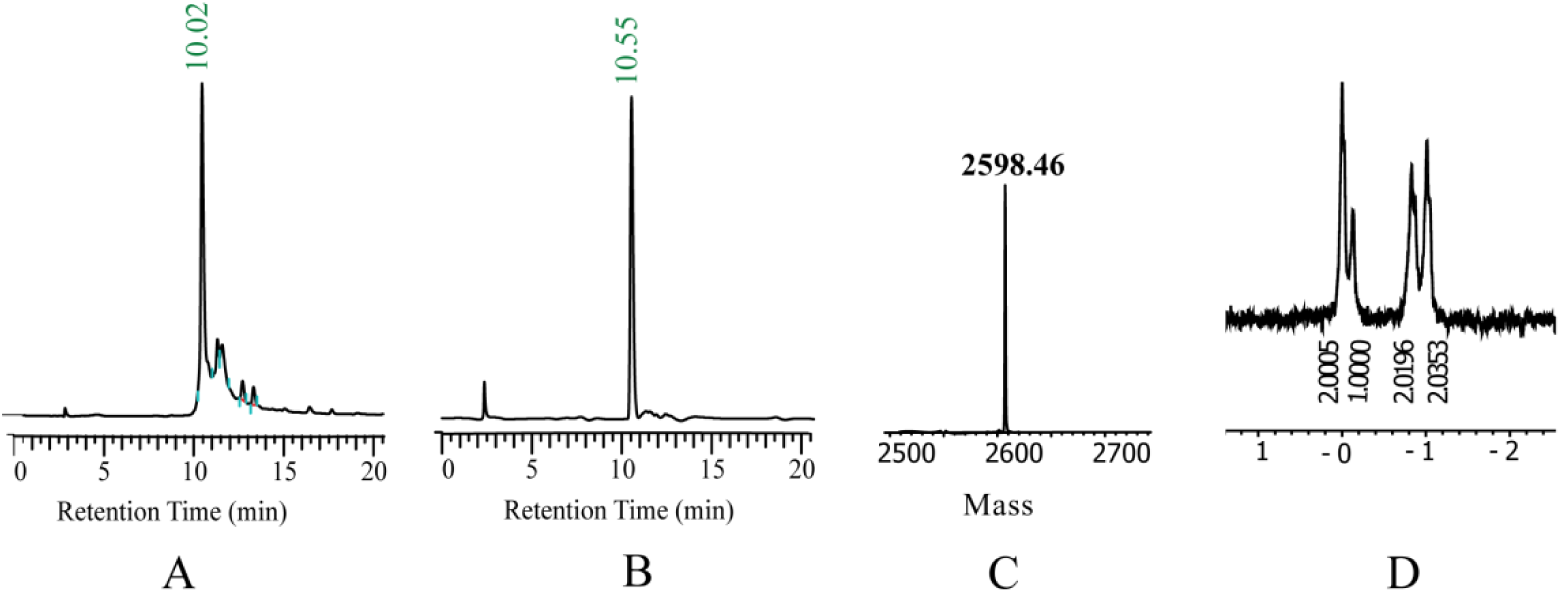
(A) RP-HPLC chromatogram of the crude seven phosphorylated peptide **1** synthesized employing our new combinatorial coupling method strategy; (B) chromatogram of the pure peptide **1**; (C) ESI-MS of the pure peptide **1**; (D) ^31^P NMR of peptide **1**.

To further improve the developed strategy, a series of multi-phosphorylated peptides **2-11** derived from Rho(330-348) with various phosphorylation patterns of 1-, 3-, 4-, 5-, 6- and 7-phosphorylations at different sites, were synthesized using the above described combinatorial approach.

Similar to peptide **1**, the coupling methods were changed gradually based on analysis of each coupling step. The coupling conditions required for the introduction of each pAA depended on a combination of factors, including the position of the pAA, whether the pAA is pSer or pThr, the proximity of the pAAs to each other and the entire phosphorylation pattern. All the phosphorylated peptides **2-11** were obtained with high purity using our combinatorial approach (Table 2). The ^31^P NMR data of the synthesized multi-phosphorylated peptides exactly matches with the number of phosphorylation sites in each peptide (Table 2). For full synthesis details and characterization data of all the peptides see Fig S3-S43.

**Table 2:** The library of phosphorylated peptides with different phosphorylation patterns synthesized in this study ^*^Details of Mass analysis data, crude RP-HPLC purity and the number phosphorylation residues determined by ^31^P NMR were provided.

| Peptide | Sequence | Observed mass (Da) | Calculated mass (Da) | Crude HPLC purity | No. of p-residues determined by $^{31}\text{P}$-NMR |
| --- | --- | --- | --- | --- | --- |
| **1** | CDDEApSpTpTVpSKpTEpTpSQVAPA | 2598.46 | 2597.90 | 76 | 7 |
| **2** | pSKpTEpTpSQVAPA | 1436.84 | 1436.56 | 90 | 4 |
| **3** | DDEApSpTpTVpSKpTEpTpSQVAPA | 2496.22 | 2495.66 | 75 | 7 |
| **4** | DDEASTTVpSKTETSQVA | 2015.51 | 2015.86 | 92 | 1 |
| **5** | DDEASpTpTVpSKTETSQVAPA | 2174.87 | 2175.79 | 87 | 3 |
| **6** | CDDEASpTpTVSKTETpSQVAPA | 2278.44 | 2279.90 | 89 | 3 |
| **7** | DDEApSTpTVSKpTETpSQVAPA | 2255.63 | 2255.76 | 87 | 4 |
| **8** | ASpTpTVpSKpTEpTSQVAPA | 1976.35 | 1976.62 | 82 | 5 |
| **9** | CDDEApSpTpTVpSKpTETSQVAPA | 2438.65 | 2437.90 | 80 | 5 |
| **10** | CDDEASpTpTVpSKpTEpTpSQVAPA | 2518.56 | 2517.90 | 78 | 6 |
| **11** | pTpTVpSKpTEpTpSQVAPA | 1897.70 | 1898.52 | 75 | 6 |

\*Details of Mass analysis data, crude RP-HPLC purity and the number phosphorylation residues determined by $^{31}\text{P}$ NMR were provided. .

Based on the synthesis of the library, we reached conclusions that can be used as general guidelines for the synthesis of other multi phosphorylated peptides. The coupling of amino acids up to first pAA can be performed using standard MW coupling method (MW-SPPS) using 5 equiv. of amino acids and Fmoc removal by MW irradiation (Table 1). The first two C-terminal pAAs could be inserted using pSPPS coupling method utilizing only three equivalents of pAAs irrespective of their type, location and proximity and removing Fmoc without irradiation (pSPPS, Table 1). Thus, this is a general trend that does not depend on the pattern, the type and the proximity of pAAs. The third pAA could also be introduced successfully using only three equivalents of pAA (pSPPS). However, increasing the coupling time from five to ten minutes (extended time phosphorylated amino acids method ET-pSPPS, Table 1) was required. All the other coupling parameters remained the same. Thus, for up to three pAAs, only three equivalents of pAAs were sufficient for achieving a complete coupling (ET-pSPPS method).

The introduction of the fourth pAA was pattern dependent. While in some cases it could be efficiently coupled employing the ET-pSPPS method (Table 1&2 peptides **1-3**, **7**, **10**, **11**), in other peptides these conditions led to the incomplete couplings (Table 1 peptides **8** and **9**). In all cases where pSer was the fourth pAA, using ET-pSPPS cycle was efficient. However, when the forth pAA was a pThr, using the ET-pSPPS cycle was not efficient enough due to the steric hindrance that slows down the coupling of Fmoc-Thr(HPO_3_Bzl)-OH. In this case, two coupling cycles were used employing 3 equiv of pAAs and 10 min coupling (double coupling pSPPS method, DC-pSPPS Table 1). The DC-pSPPS coupling method was also efficient for the introduction of the fifth pAA except for peptide **8**. The introduction of the fifth pAA in peptide **8**, similar to the introduction of the sixth pAA in peptides **1**,**3**,**10**, and **11** was extremely difficult because Fmoc-Thr(HPO_3_Bzl)-OH was coupled directly to an adjacent protected phosphorylated Thr residue and peptide **8** already contains three pThr residues. In this case, using five equivalents of Fmoc-Thr(HPO_3_Bzl)-OH and repeating the reaction cycle provided the suitable method for the insertion of the fifth pAA (excess building block pSPPS method, EBB-pSPPS, Table 1). These conditions were also essential for the introduction of the sixth and seventh pAAs in all remaining peptides (Table 1 peptides **1**, **3**, **10**, **11**).

Using EBB-pSPPS cycle proved excellent for the synthesis of even the most hindered peptide motifs. The only drawback of this coupling method is the requirement of ten equivalents of each pAA. However, applying this EBB-pSPPS coupling cycle for the insertion of all pAAs in peptide **1** would require the overall use of 70 equivalents of these highly expensive pAAs. By using the developed combinatorial strategy the overall amount of pAA used for the synthesis of peptide **1** is almost half.

## Conclusions

In conclusion, our study presents an efficient combinatorial approach for the preparation of multi-phosphorylated peptides. By understanding the combination of factors that influence the coupling yields of each pAA, we were able to develop a cost effective yet high yielding synthetic strategy even to the most difficult-to-synthesize multi-phosphorylated peptides. This method can be used as a guideline for the design and synthesis of libraries of specific peptides with different phosphorylation patterns. The accessibility to these extremely important entities is essential for future research in many areas of chemical biology and drug design.

## Acknowledgements

AF is supported by grants from the Israel Science Foundation and the Israel Cancer Research Foundation and by the Minerva Centre for Bio Hybrid complex systems. MH was supported by the EU project RECORD-IT-AMD-664786. S.M is thankful to Lady Davis fellowship trust of Hebrew University for post-doctoral fellowship. DBV and DM were supported by the Swiss National Science Foundation grants 141898, 159748 to DBV. We thank Luxembourg industries limited, Israel for their assistance.

## Associated content

Supporting information of this article contains the following details: Additional figures, synthesis procedures and characterization data of all the peptides, copies of ESI-MS, UPLC and ^31^P NMR.

